# Bread and hummus: trait connectance and correlation pleiades in grain crops

**DOI:** 10.1101/2024.05.22.595267

**Authors:** Victor O. Sadras

## Abstract

Phenotypic integration has been investigated from multiple perspectives. From a developmental perspective, connectance has been defined as the level of linkage between traits. Correlation pleiades, *i.e*., correlations between some traits and, simultaneously, lack of correlations between these and other traits have been interpreted as the independence of certain developmental processes with respect to other processes within the organism, and as the outcome from the discrepancy between the agencies participating in the formation of the trait and the selective forces influencing its function. Here, I use two published data sets to test the variation in connectance with both trait and genotype and the existence and meaning of correlation pleiades in wheat and chickpea. Connectance varied from 0.09 to 4.2 in wheat and from 0.06 to 22.8 in chickpea, and cluster analyses revealed correlation pleiades. The frequency distribution of connectance conformed to a power law with similar slopes = −1.665 ± 0.222 for wheat and −1.555 ± 0.126 for chickpea, consistent with developmental self-organisation. Connectance was lower for traits with higher heritability such as seed weight, which together with the negative association between heritability and phenotypic plasticity completes a relational triangle: high connectance ⇔ low heritability ⇔ high phenotypic plasticity.

## Introduction

Connectivity and phenotypic integration have been investigated from evolutionary, ecological, developmental and breeding perspectives (Amzallag, 1999a, b, 2001, 2004; Armbruster *et al*., 1999; Berg, 1960; Climent *et al*., 2024; Damián *et al*., 2018; Damián *et al*., 2020; Leslie and Mander, 2023; Pigliucci, 2003). Except for Amzallag’s work with sorghum, most of these studies looked at wild plants and model species. Phenotypic integration in grain crops grown under field conditions, the focus of this article, has been overlooked.

The concept of phenotypic integration was incipient in the 1960s but has been in the background of evolutionary and ecological research throughout the second half of the 20th century partly because of the shift to molecular techniques “that have focused attention elsewhere and partly because longstanding conceptual and analytical problems have remained” (Pigliucci, 2003). On evolutionary time scales, major reproductive innovations like the origin of seeds and angiospermy correlated with increased phenotypic integration quantified as interactions among plant parts in network analysis (Leslie and Mander, 2023). However, caution is needed in interpreting these correlations as causal relationships because for certain groups, particularly Mesozoic gymnosperms, reproductive innovations emerged much earlier than increased interactions (Leslie and Mander, 2023).

From a developmental perspective, connectance has been defined as the level of linkage between traits (Amzallag, 2000). Development is not a continuous process but alternates stabilisation during discrete stages, *i.e*., phenophases, and dismantling of relational networks between organs in the intervening, shorter critical periods (Amzallag, 2004). In this context, the critical period is a transient phase of isolation of the system that enables its evolution towards equilibrium; the transition from dissipative to isolated system is the source of newly emerging dissipative structures, *i.e*., new phenophases, in which environmental or developmental disturbances are adaptively integrated (Amzallag, 2004). This developmental definition of “critical period” should not be confounded with the agronomic definition of “critical period” where crop yield is more sensitive to stress (Carrera *et al*., 2024). Consistently with Amzallag’s (2000, 2004) propositions, integration of foliar traits in Turnera velutina, an endemic Mexican shrub, increased from juvenile plants, which featured two functional modules related to plant defence and leaf economy, to reproductive plants with greater interconnectivity and hence lower modularity. The ontogenetic changes in foliar trait integration were considered adaptive by allowing plants to accommodate changing selective dynamics and physiological priorities through development (Damián *et al*., 2018).

From an eco-physiological perspective, correlations are expected between traits involved in resource acquisition, defence, and stress tolerance due to resource trade-offs, multifunctionality of traits and/or regulatory processes. Two examples in this context. Damián *et al*. (2020) estimated intraspecific variation in leaf functional traits related to the primary metabolism and anti-herbivore defence in Turnera velutina. Phenotypic integration of leaf traits varied 10-fold among 13 maternal families and correlated with relative growth rate and production of flowers, but not with production of seed. Amzallag (1999a) showed that exposure to a sublethal concentration of NaCl during early vegetative development increased tolerance to salinity in sorghum. The phase of competence for induction of this response coincides with the emergence of the first adventitious roots and is higher in genotypes with a weaker link between the seminal root and the shoot during the emergence of the adventitious root. This led to the conclusions that: (i) functional integration of the adventitious roots within the whole plant has an adaptive nature in normal development; (ii) salt adaptation results from an integration of the environmental constraint (NaCl) during this developmental readjustment; and (iii) perturbations associated with the emergence of a new organ cause rapid variations in sensitivity required to open a competence window. In both cases, the lens of phenotypic integration has been insightful.

Correlation pleiades, *i*.*e*., correlations between some traits and, simultaneously, lack of correlations between these and other traits have been interpreted from developmental and evolutionary perspectives (i) as the independence of certain developmental processes with respect to other processes within the organism, and (ii) as the outcome from the “discrepancy between the agencies participating in the formation of the character and the selective forces determining its function” (Berg, 1960). For example, correlation pleiades were apparent in plants relying on specialised pollinators were vegetative traits correlate among themselves but not with flower traits (*e*.*g*., *Genarium platense, Linaria vulgaris*) but not in selfers (*e*.*g*., *Triticum eaestivum, Hordeum vulgare*) or wind-pollinated plants (e.g., *Elimus arenarius*) where correlations between vegetative and reproductive traits are stronger (Berg, 1960). Further tests of predictions from Berg’s (1960) hypothesis returned mixed results, and supported the conclusion that the patterns of integration among floral traits and between floral and vegetative traits are species specific rather than a consequence of pollination ecology (Armbruster *et al*., 1999).

In this article, we test the variation in connectance with both trait and genotype and the existence and meaning of correlation pleiades in field-grown wheat and chickpea.

### Data and calculations

We analysed two published data sets including 17 traits measured in 13 wheat varieties grown in four environments (Sadras and Lawson, 2013) and 20 traits in 20 chickpea varieties grown in eight environments (Sadras *et al*., 2016). Connectance was calculated in two steps (Amzallag, 2000). First, correlation coefficients *r* were *z*-transformed to account for departure from normal distribution (eq. 1), and connectance *C*(*T*_*k*_) for each trait *T* was calculated as the average of the module of *z* (eq. 2). Correlation pleiades were established with cluster analysis of connectance using the Ward method standardised by column (JMP version 17.2.0).

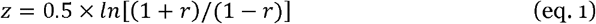

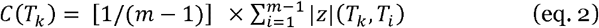

### The frequency distribution of connectance conformed to a power law in wheat and chickpea data sets

Connectance varied from 0.09 to 4.2 in wheat and from 0.06 to 22.8 in chickpea. In both crops, the frequency distribution of connectance conformed to a power law with slope = −1.665 ± 0.222 for wheat and −1.555 ± 0.126 for chickpea (Fig. 1ab). Genotype-dependent median connectance in wheat varied from 0.28 in Condor to 0.67 in Gladius (Fig. 2a). In this collection of genotypes, Condor was the first semi-dwarf cultivar and Gladius was the newest cultivar (Fig. 2a). The introduction of *Rht* genes associated with a dramatic reduction in connectance for Condor, with further breeding and selection restoring connectance (Fig. 2a). Connectance of yield increased at 0.028 per year over the period 1958 to 2007 (inset Fig. 2a). The genotype-dependent median connectance in chickpea varied from 0.29 to 0.77 (Fig. 2b). In comparison to the 2.4-fold variation in wheat and 2.7-variation in chickpea, intraspecific variation in phenotypic integration varied 10-fold in a population of *Turnera velutina* (Damián *et al*., 2020).

**Fig. 1.**
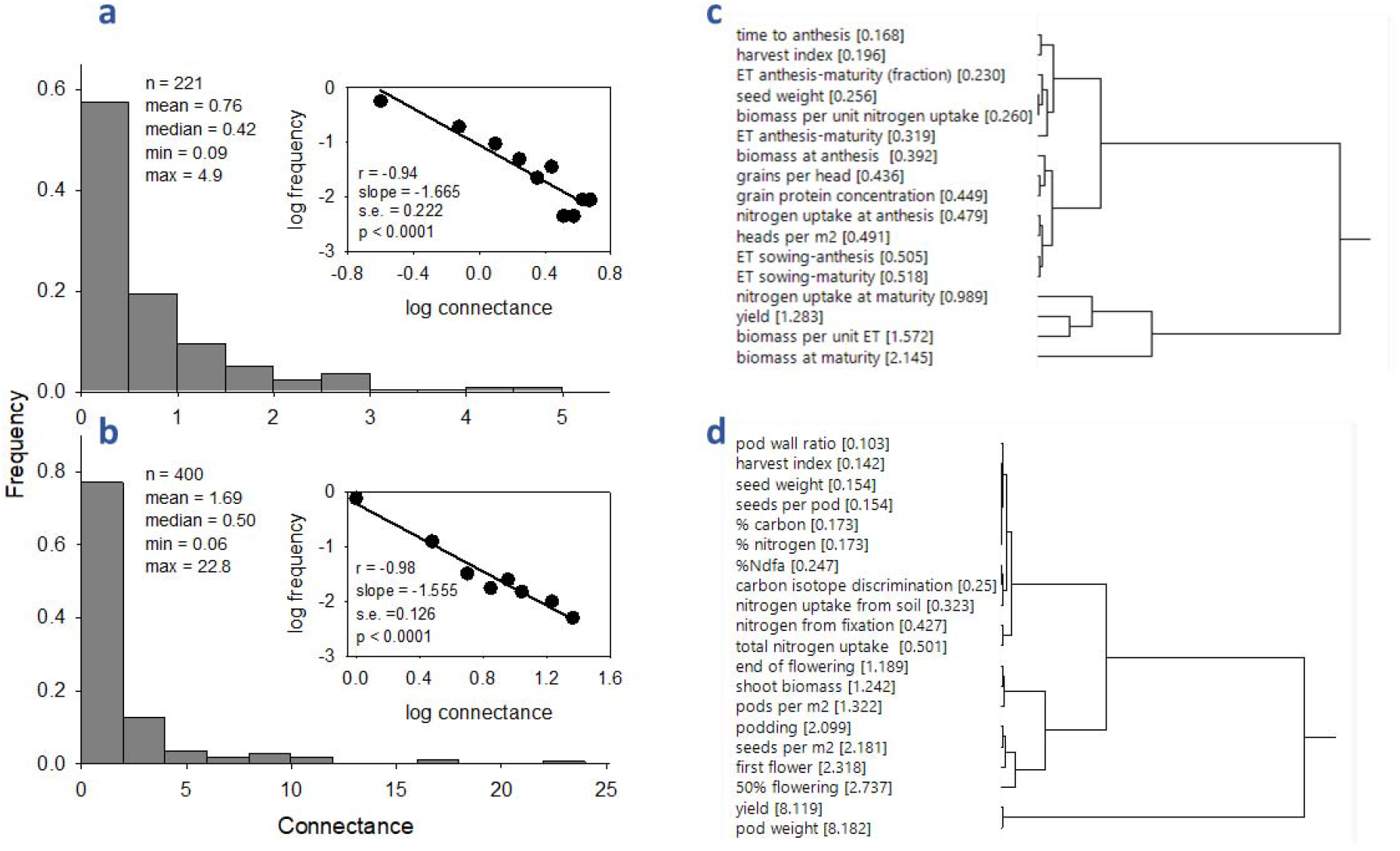
(a, b) Frequency distribution of connectance and (c, d) hierarchical clustering of connectance for traits in (a, c) wheat (b, d) chickpea. Data comprise 17 traits measured in 13 genotypes for wheat (n = 221), and 20 traits measured in 20 genotypes for chickpea (n = 400). Wheat was grown in four field environments and chickpea in eight field environments. Insets show frequency distributions in a log-log scale where the line is the least squares regression. Clusters calculated with Ward method. Data sources: Sadras et al. (2013) for wheat and Sadras et al. (2016) for chickpea.

**Figure 2.**
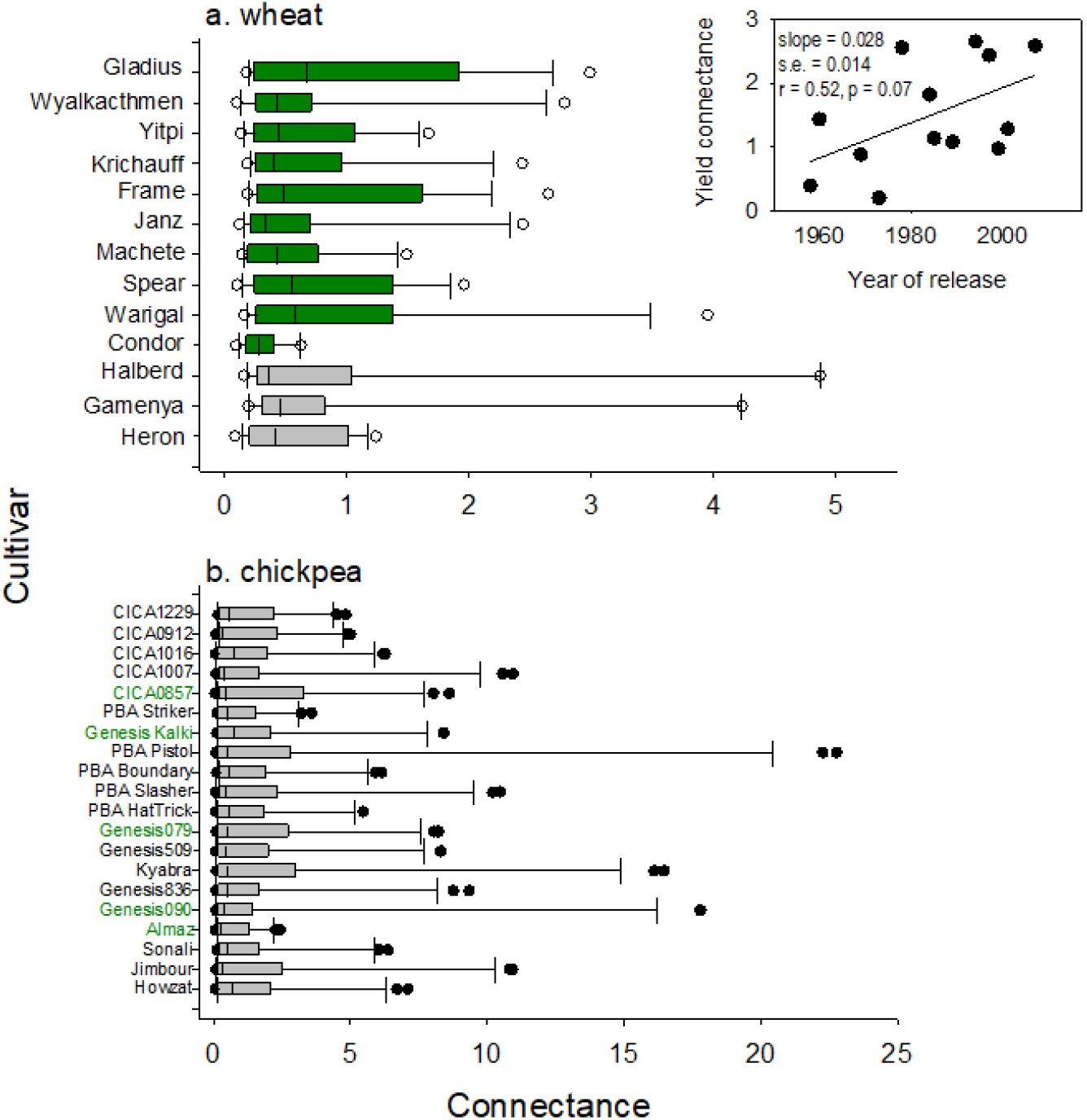
Genotype-dependent variation in connectance of (a) 17 traits in wheat and (b) 20 traits in chickpea. Genotypes are ordered by year of release, from 1958 (bottom) to 2007 (top) in wheat, and from 2001 (bottom) to 2017 (top) in chickpea. Inset is the variation in connectance of wheat yield with year of cultivar release between 1958 and 2007. For wheat, grey is tall, pre-green revolution cultivars, and green are semi-dwarf. For chickpea, green labels are Kabuli and black labels are Desi. Data sources: Sadras et al. (2013) for wheat and Sadras et al. (2016) for chickpea.

Diverse systems, including genetic and metabolic networks, feature complex topology with a common property: the vertex connectivities follow a scale-free power-law distribution (Bak, 1996; Barabási and Albert, 1999; Kauffman, 1995). This feature is a consequence of two generic mechanisms: (i) networks expand continuously by the addition of new vertices, and (ii) new vertices attach preferentially to sites that are already well connected. A model based on these two mechanisms reproduced scale-free distributions, which indicates that the development of large networks is governed by robust self-organising phenomena that go beyond the particulars of the individual systems (Barabási and Albert, 1999; Kauffman, 1995). Power-law distributions are consistent with but do not prove self-organisation.

With a focus on connectance, Amzallag (1999b) advanced a biological explanation for a role of self-organisation in developmental process. Experiments with sorghum showed a negative relationship between connectance and heritability calculated from parent-offspring relationships. For both wheat and chickpea, connectance was low for seed weight (Fig. 1cd) in alignment with the high heritability of this trait, usually above 0.7 (Harper *et al*., 1970; Sadras, 2007). For wheat, time to flowering had the lowest connectance of all 17 traits tested, in correspondence with the high heritability of this trait, *e*.*g*., broad sense heritability of time to heading and flowering between 0.7 and 0.9 (Dunckel *et al*., 2017). For chickpea, phenological development had high connectance, above 2 (Fig. 1d); this compares with lower heritabilities, *e*.*g*., below 0.5 for time to flowering and below 0.4 for podding from parent-offspring correlations (Anbessa *et al*., 2006). Whereas daylength and temperature are drivers of phenological development in both wheat and chickpea, soil factors such as moisture and salinity also modulate phenological development of pulses (Li *et al*., 2022; Lizarazo *et al*., 2017); a larger environmental component of the phenotypic variance can therefore account for the lower heritability of phenological development in chickpea and its associated higher connectance. In both crops, connectance of yield was at the higher end of the range (Fig. 1cd) in correspondence with the low heritability of yield for which the components of the phenotypic variance usually rank environment >> genotype x environment interaction > genotype, (Cooper *et al*., 1995; Hoffmann *et al*., 2009; Yang *et al*., 2005).

The correlation between connectance and heritability could be interpreted in two alternative ways: (1) pre-existing information, as quantified with heritability, is the single source of developmental control whereas the linkage between traits generates a “developmental noise” that disturbs the expression of this pre-existing information or (2) the network of relationships is also a source of information for development, “which completes, substitutes or even counteracts expression of pre-existing information” (Amzallag). Contradictory evidence, which supported both (1) and (2), was interpreted with a speculative model based on the variability of connectance whereby (1) applies above and (2) applies below a threshold of connectance: “below a critical value of lability, expression of a character is controlled by its connectance… and… an increased lability in connectance reduced its involvement in character expression, which became probably mainly determined by a pre-existing information” (Amzallag, 1999b). Further considerations on cell-to-cell signalling during development lead to the conclusion that “the metacellular network is not directly determined by pre-existing information but is an expression of the self-organization dimension of development. Thus, although related to genetic expression, connectance may be considered as an autonomous dimension in development.” Amzallag (1999b) conclusion that morphogenesis involves both genetic and self-organising dimensions, has implications for the reliability, stability and adaptability to the developmental processes. From a teleonomic (*i*.*e*., purpose oriented) perspective of development, Levin (2023) also concludes that “development is incredibly reliable, producing bodies to very tight tolerances despite considerable deviations and noise at the level of gene expression and cellular activity” … “development is not hardwired but context-sensitive and plastic”.

The emphasis on “data sets” in the title of this section is to caution against generalisations; the findings reported here are specific for the combinations of crop species, traits, and growing conditions; other data sets may feature different frequency distributions, *e*.*g*., log-normal (Broido and Clauset, 2019). A small sample from an experiment comprising eight traits measured in six sorghum lines grown in a single environment (n = 48), returned a close-to-normal frequency distribution of connectance (Shapiro-Wilk test p = 0.06), with a range from 0.17 to 1.14, median = 0.63 and mean = 0.60 (calculated from data in Table 2 of Amzallag, 1999b).

### Correlation pleiades were apparent for both wheat and chickpea data sets

Correlations between traits are commonplace in studies of plant and crop phenotypes. Positive correlations do not prove but are a first step towards demonstration of causal relations (e.g., Calderini *et al*., 2021). Negative correlations are mostly interpreted in terms of trade-offs (e.g., Henery and Westoby, 2001). Beyond these two general angles, interpretation of correlations usually lacks deeper theoretical context.

Berg (1960) influential notion of correlation pleiades allows for developmental and evolutionary interpretations of plant phenotypes. From this lens, Berg (1960) concludes that “correlation coefficients between the dimensions of grains and those of other parts of the organisms are somewhat smaller than all other correlation coefficients. This might well be expected, the grain being essentially an organism of the next generation with a genotype of its own, and not a part of the parent organism.” The typically high heritability of grain weight and grain dimensions has been interpreted in terms of Smith and Fretwell’ s (1974) mother-offspring conflict (Sadras, 2007). The low connectance of seed weight in both wheat and chickpea (Fig. 1cd), and the negative association between heritability and phenotypic plasticity (Alvarez Prado *et al*., 2014; Lacaze *et al*., 2009) completes a relational triangle: high connectance ⇔ low heritability ⇔ high phenotypic plasticity.

## Conclusion

Connectivity is a central theme for networks at small, *i.e*., molecular, and high, *i.e*., ecosystem scales. At the scale of the individual plant, connectivity has received less attention and is not part of our current thinking in crop sciences. We propose that studying the genotype- and trait-dependent variation in phenotypic integration could be useful to understand and manipulate plant development against the relational triangle with connectance, heritability and plasticity vertices.

